# Fundamental and realized feeding niche breadths of sexual and asexual stick-insects

**DOI:** 10.1101/377176

**Authors:** Larose Chloé, Darren J. Parker, Schwander Tanja

## Abstract

The factors contributing to the maintenance of sex over asexuality in natural populations remain largely unknown. Ecological divergences between lineages with different reproductive modes could help to maintain reproductive polymorphisms, at least transiently, but there is little empirical information on the consequences of asexuality for the evolution of ecological niches. Here, we investigated how niche breadths evolve following transitions from sexual reproduction to asexuality. We estimated and compared the realized feeding niche breadths of five independently derived asexual *Timema* stick insect species and their sexual relatives. We found that asexual species had a systematically narrower realized niche than sexual species. To investigate how the narrower realized niches of asexual versus sexual species come about, we quantified the breadth of their fundamental niches but found no systematic differences between reproductive modes. The narrow realized niches found in asexuals are therefore likely a consequence of biotic interactions that constrain realized niche size in asexuals more strongly than in sexuals. Interestingly, the fundamental niche was broader in the oldest asexual species compared to its sexual relative. This broad ecological tolerance may help explain how this species has persisted over more than a million years in absence of sex.

## INTRODUCTION

The maintenance of obligate sex in natural populations, despite numerous disadvantages compared to other reproductive systems, is a major evolutionary paradox. Although there is a rich body of theory proposing potential benefits of sex, empirical studies evaluating such benefits under natural conditions remain scarce (reviewed in Neiman et al. 2018). A simple mechanism that could facilitate the maintenance of reproductive polymorphisms is niche differentiation between sexual and asexual species (Meirmans et al. 2012). Such niche differentiation could result from a difference in ecological optima between sexuals and asexuals (e.g., Case and Taper 1986), or from situations where sexual species cover larger fractions of the available niche space than their asexual counterparts (e.g., Bell 1982).

Because asexual species derive from sexual ancestors, fundamental niches (i.e., the range of environmental conditions that allow for survival, growth and reproduction) in new asexual species should depend directly upon the fundamental niche found in the ancestral sexual species. How the fundamental niche in an ancestral sexual population translates to that found in an asexual population is however unclear. For example, the *Frozen Niche Variation* model (FNV) predicts that the phenotypic distribution of a new, recently derived asexual would be narrower than that of its genetically variable sexual ancestor, because a single sexual genotype will be “frozen” in the new asexual lineage (Vrijenhoek 1984, Case and Taper 1986, Case 1990, Weeks 1993, Vrijenhoek and Parker Jr 2009; Fig, 1A). By contrast, the *“General-Purpose Genotype*’ hypothesis (GPG; Lynch 1984; but see also Baker 1965; Parker et al. 1977) proposes that asexual lineages should generally have broader environmental tolerances than sexual individuals because of strong selection for plasticity in asexuals. Under this scenario, we would expect asexual populations to have broader ecological niches than sexual ones (Fig. 1B). The two hypotheses are non-mutually exclusive. For example, by combining the FNV and GPG, we can suggest that young asexual lineages would feature, on average, narrow niches, while old ones would feature broad niches.

**Figure 1.**
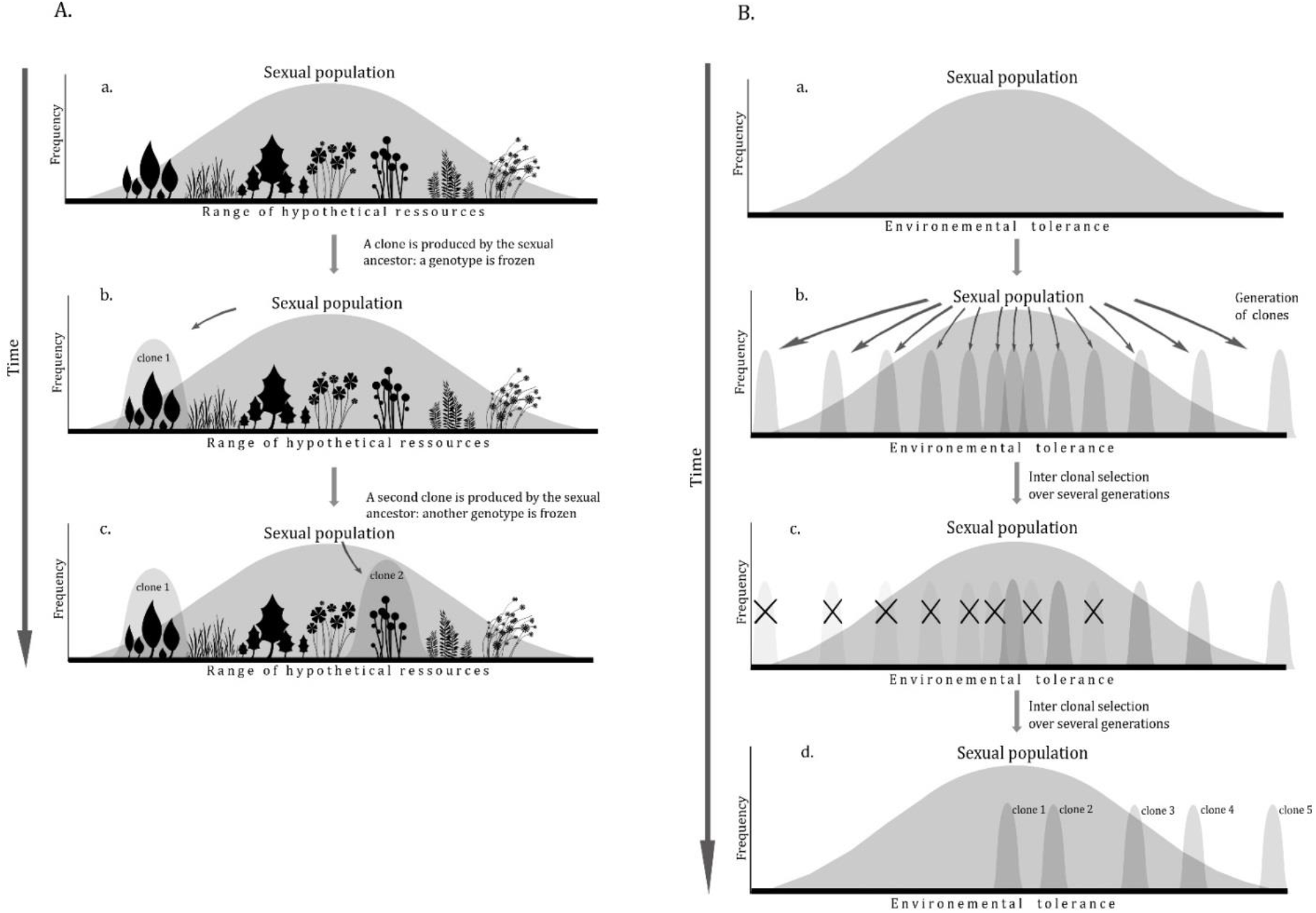
**(A) The frozen niche variation model.** (a) A sexual population (broad curve) exhibits genetic variation for the use of a natural resource (here symbolized by a range of hypothetical plants). (b) A new asexual clone is produced, comprising a small subset of the genotypic diversity contained in its sexual ancestor (c) A second clone is produced from a different sexual genotype characterized by a different ecological niche. The niche breadth of the sexual population as a whole is larger than the one of each individual clone. Figure modified from Vrijenhoek and Parker, 2009. **(B) General purpose genotype hypothesis.** (a) Individuals in a sexual population vary in the range of their environmental tolerances (narrow to broad phenotypic plasticity) (b) Clones are produced from different genotypes in the sexual population with different levels of phenotypic plasticity. (c and d) Natural selection favors clones with broader tolerances such that clones may feature higher levels of phenotypic plasticity than the sexual population as a whole (e.g. extreme case of clone 5). Figure adapted from Vrijenhoek and Parker, 2009.

Regarding the breadth of the realized niche (i.e., the fraction of the fundamental niche used by organisms under natural conditions), there is currently no specific theory predicting similarities or differences between sexuals and asexuals. There are however several theories predicting that sex can accelerate the rate of adaptation compared to asexuality (Hill and Robertson 1966, Kondrashov 1988, Barton and Charlesworth 1998, Otto and Lenormand 2002). Sexual organisms therefore may be able to evolve adaptations to competitors, pathogens, or predators more rapidly than asexuals. As a consequence, the realized niche in asexual organisms may be smaller than in sexual organisms due to a reduced ability to respond to these biotic pressures.

Here we evaluate whether asexuality is associated with different niches and niche sizes than sexual reproduction, using herbivorous stick insects of the genus *Timema* as a model system and different host plants as a proxy for different niches. Seven independently derived asexual lineages have been identified in this genus, each with a closely related sexual counterpart (Schwander et al. 2011; Fig. 2). This allows us to perform replicate comparisons between sexual and asexual lineages. Moreover, the asexual *Timema* lineages vary in age (Law and Crespi 2002, Schwander et al. 2011), allowing us to assess the possible consequences of asexuality on niche breadth over a range from recently derived to long-term asexuality.

**Figure 2.**
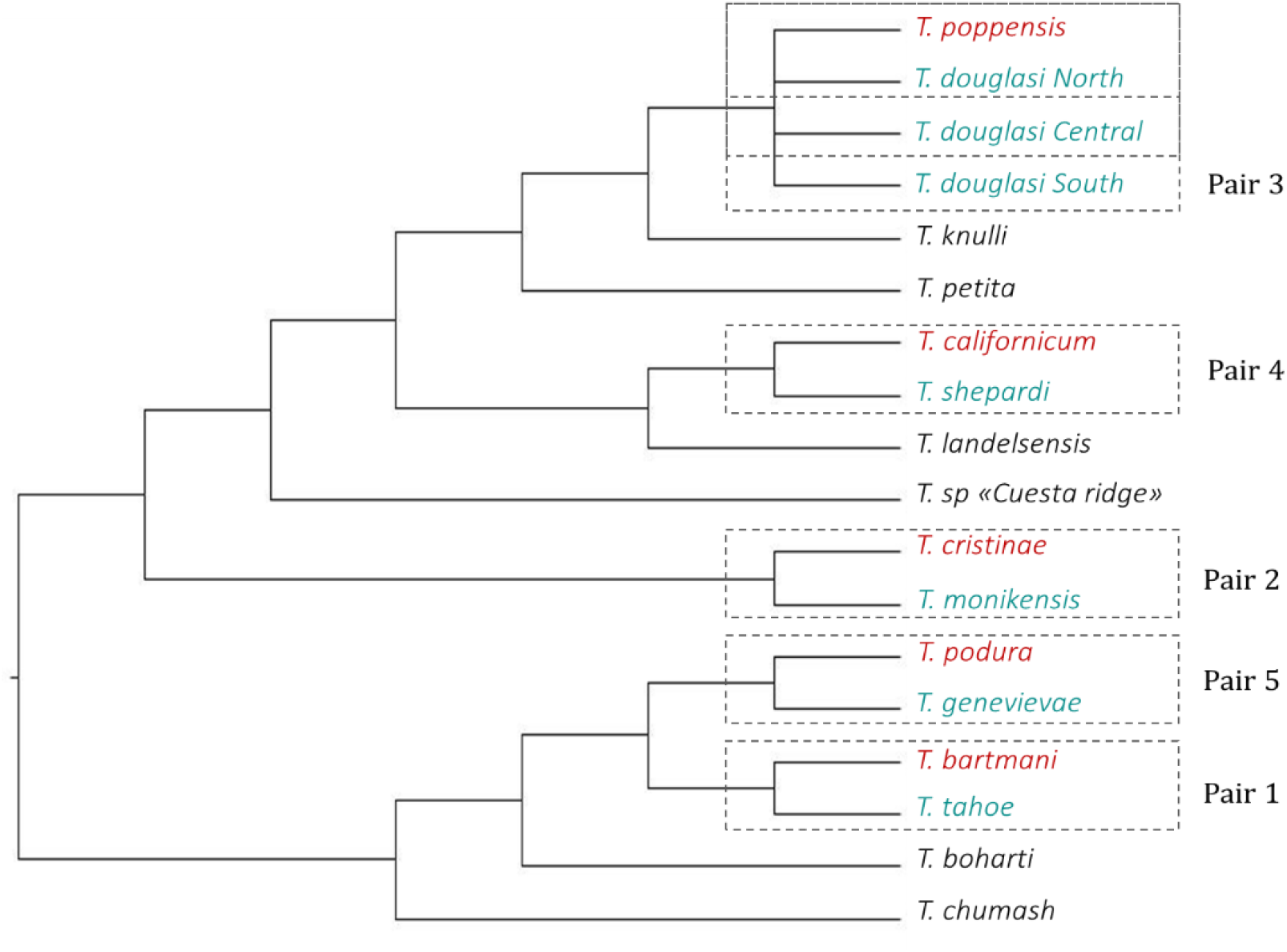
*Timema* phylogeny highlighting the sexual and asexual species. Phylogeny redrawn from Riesch *et al*. (2017) with the seven asexual lineages added from Schwander *et al*. 2011 (in blue). The used sexual species are labeled in red. Pair numbers correspond to the most recent (i.e., pair 1) to the most ancient (i.e., pair 5) transition to asexuality (ranking from Bast *et al*. 2018).

We first estimated the size of the realized feeding niches of sexuals and asexuals both at the species and at the population level in five sexual-asexual *Timema* sister species pairs, using occurrences on different host plants in natural populations. *Timema* feed on the leaves or the needles of very diverse host plants, comprising both angiosperms and conifers, and the quality of these plants as a food source is highly variable (Larose et al. 2018). We then conducted feeding experiments with species from four sexual-asexual species pairs to estimate the size of their fundamental feeding niches. Finally, we evaluated the contribution of predation to shaping realized niches in sexuals and asexuals. *Timema* are characterized by different cryptic morphs on different host plants, both within and between species (Sandoval 1994a, 1994b, Nosil 2007, Sandoval and Crespi 2008). Previous studies have shown that the combination of selection imposed by predators and *Timema* host preference maintain a correlation between morph frequency and host-plant frequency between populations (Sandoval 1994a, Nosil 2004, Sandoval and Nosil 2005), indicating that color polymorphism and predation may be of key importance for realized niches in *Timema*.

## METHODS

### Realized feeding niche breadths

Data from a previous study that collected information on host plant use across all 23 known *Timema* species (Larose et al. 2018) allowed us to estimate the size of the realized feeding niche of sexuals and asexuals at the species level. To estimate the realized niche at the population level, we further performed a count of the number of individuals collected on each potential host plant across 30 populations from five species pairs (between two and six populations per species; Table S2). The size of the realized feeding niche per population was then quantified with the inversed Tau (τ) specialization index (Yanai et al. 2004), which ranges from 0 (pure specialist) to 1 (complete generalist).

### Degree of color polymorphism

Color phenotypes vary broadly in several *Timema* species but can be separated into a total of 14 discrete morphs across all species (range 1-8 per species; Table S1). We recorded color morph frequencies from all sampling locations (Table S2) and used the Simpson diversity index to quantify the level of polymorphism (Simpson 1949). This index varies between 0 (here indicating color monomorphism) and 1 (indicating diversity of equally frequent color morphs). We then estimated the correlation between the degree of color polymorphism and the size of the realized feeding niche, both at the species and at the population levels with Phylogenetic Generalized Least Squares (PGLS) to account for phylogenetic non-independence among *Timema* species. These analyses were conducted using the ape (Paradis et al. 2004) and nlme (Pinheiro et al. 2009) R packages (R Core Team 2017) using a Brownian motion model for trait evolution.

### Fundamental feeding niche breadths

To estimate the fundamental feeding niche breadths of sexual and asexual *Timema* species, we performed a feeding experiment and measured insect performance on different host plants. We chose seven plants known to be commonly used by several *Timema* species, while trying to cover the phylogenetic diversity of the host plants (Larose et al. 2018). Specifically, we chose four angiosperms: (*Ceanothus thyrsiflorus* (lilac, lil), *Adenostoma fasciculatum* (chamise, cha), *Quercus agrifolia* (oak), and *Arctostaphylos glauca* (manzanita, mz)), and three conifers: (*Pseudotsuga menziesii* (douglas fir, df), *Abies concolor* (white fir, wf), and *Sequoia sempervirens* (redwood, rdw)). Stick insects from eight *Timema* species (four sexual-asexual species pairs) were collected from multiple field sites in California (Table S3). We only used fourth-instar juvenile females for feeding experiments to minimize age-related effects on insect performance during our experiments. Between 10 and 20 such females were used per host plant to measure survival and weight gain during 10 days, for a total of 70-105 females per population (635 insects in total; Table S3).

We first used a generalized linear model (GLM) with a binomial error to compare survival and an ANOVA to compare the weight gain of all stick insects species on the different plants using R (R Core Team 2017). We then compared for each *Timema* species pair separately, the survival and weight gain of the sexual and asexual individuals, testing specifically for an interaction between reproductive mode and plant species, because a significant interaction between these two factors would indicate a difference in fundamental niche between sexuals and asexuals. Finally, we quantified the breadth of the fundamental feeding niche of the eight *Timema* species using again the inversed Tau index. We could not compare the fundamental niche of the *T. bartmani/T. tahoe* species pair because *T. tahoe* individuals of the appropriate developmental stage could not be collected in sufficient numbers for the feeding experiment.

## RESULTS

### Realized feeding niche breadths

For realized niches measured at the species level, the sexuals are more ecologically generalist in four out of five cases, as they used at least twice as many plants as their asexual relatives (Fig. 3A). In the remaining case (*T. poppensis/T. douglasi*), the sexual and the asexual species used the same number of host plants in the wild (Fig. 3A). For realized niches measured at the population level, all ten species are relatively specialist (Tau indices varying between 0 and 0.48; Fig. S1B) and there were no significant differences in the degree of specialization between sexual and asexual populations (GLM; p-value = 0.19). However, we did find that (within species) sexual populations vary more than asexual ones in their degree of specialization (Levene’s test, F_1, 27_ = 12.2, p-value < 0.002; Fig. S1B).

**Figure 3.**
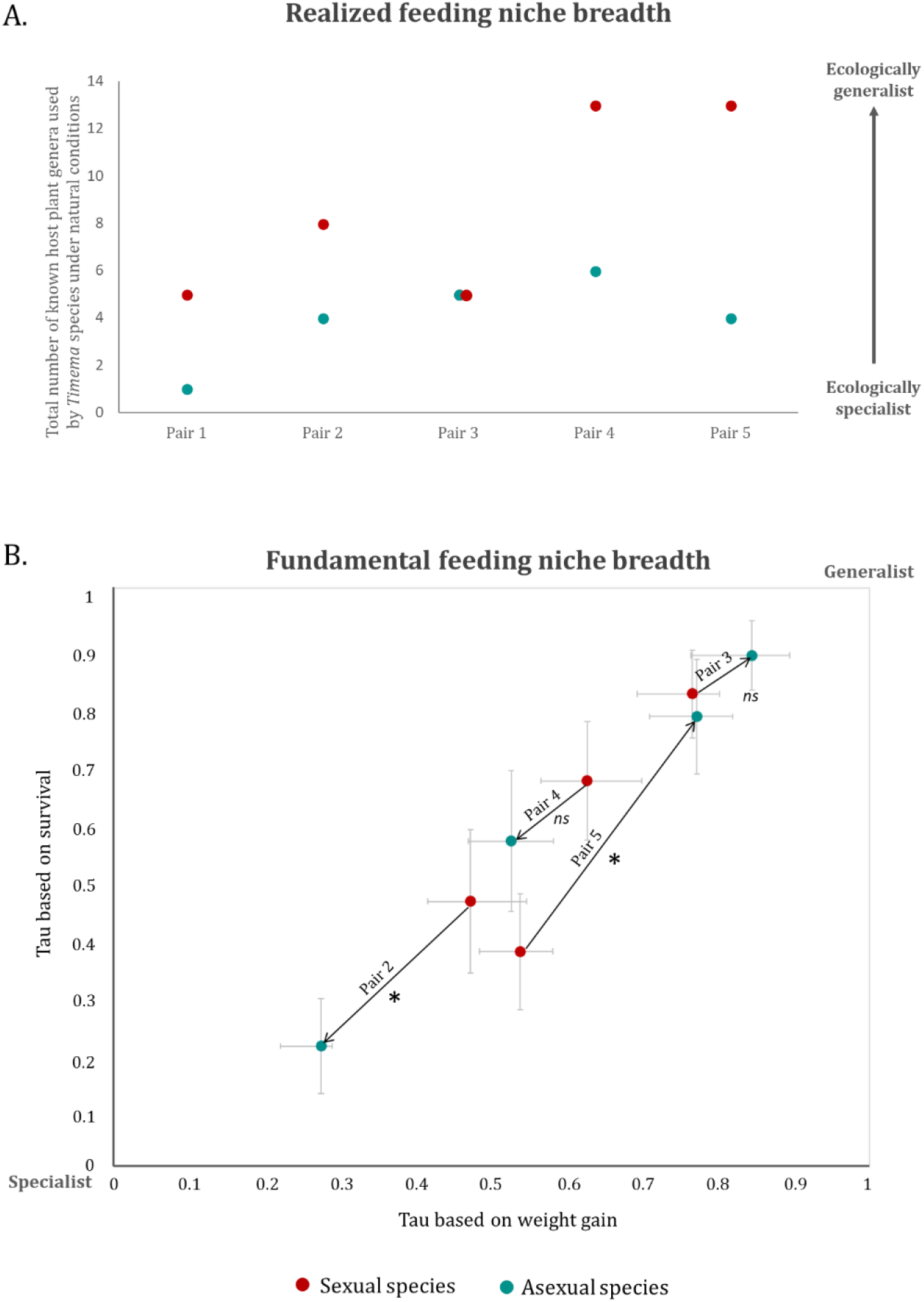
Realized (A) and fundamental (B) feeding niche breadths of sexual and asexual *Timema* stick insects, measured at the species level. The pairs are listed from the most recent to the most ancient transition to asexuality (ranking from Bast *et al*. 2018). For the size of the fundamental niche we used the specificity indices Tau based on weight gain and survival during ten days. Stars indicate significant differences of the Tau indices of the sexual and asexual species of a pair. For species pair numbers, see Fig. 2.

To assess potential interactions between color polymorphism and the number of different host plant species used, we compared the degree of color polymorphism within *Timema* species and populations with their degree of ecological specialization. At the species level, the size of the realized niche was correlated with the number of morphs of these species (correlation corrected with PGLS; r= 0.57, p-value < 0.003; Fig. S1). Similar to the size of the species-level realized niche, the asexuals contain two to five times fewer morphs than their sexual relatives, with the exception of *T. poppensis/T. douglasi*, in which both species have only a single morph (Table S1). By contrast, at the population level, we did not detect any link between color polymorphism and the size of the realized feeding niche (Pearson’s correlation; r = 0.14, p-value = 0.46; Fig. S2B).

### Fundamental feeding niche breadths

Survival and weight gain vary widely among the different studied *Timema* species when fed with different plants (p < 2.2 − 10^−16^ for survival and F_7, 292_ = 8.94, p < 5.5 − 10^−10^ for weight gain; Fig. S1A, Fig. 3B), and Tau indices based on survival or weight gain were strongly correlated (Pearson’s correlation, r = 0.96, p < 0.0001; Fig. 3B). We found significant differences in the fundamental niche breadths of sexuals compared to asexual species in two species pairs, (*T. cristinae/T. monikensis* and *T. podura/T. genevievae;* Fig. S1A, Fig. 3B). The remaining two pairs (*T. poppensis/T. douglasi* and *T. californicum/T. shepardi)* showed no significant difference (Fig. 3). Interestingly, *T. monikensis* and *T. genevievae*, which represent the most recent asexual lineage and oldest asexual lineage tested respectively, were characterized by an opposite result. *T. monikensis* was significantly more specialist (Tau based on weight gain = 0.27, 95% CI 0.22 – 0.29; survival = 0.21, 95% CI 0.13 - 0.29) than its sexual relative *T. cristinae* (Tau based on weight gain = 0.47, 95% CI 0.41 - 0.55; survival = 0.46, 95% CI 0.34 - 0.58; Fig. 3B). On the contrary, the ancient asexual *T. genevievae* was significantly more generalist (Tau based on weight gain = 0.77, 95% CI 0.71 - 0.82; survival = 0.78, 95% CI 0.68 - 0.88) than it sexual sister species *T. podura* (Tau based on weight gain = 0.54, 95% CI 0.48 - 0.58; survival = 0.37, 95% CI 0.27 - 0.47; Fig. 3B). Finally, we found that the fundamental feeding niche breadths were not correlated with the sizes of their realized feeding niche, neither at the species level (Pearson’s correlation; r= 0.13, p= 0.77; Fig. S1A), nor at the population level (r= -0.14, p= 0.50; Fig. S1B).

To test whether asexual and sexual species feature different fundamental feeding niches, we modeled, in each species pair, the survival and weight gain as functions of the species’ reproductive mode and of the experimental feeding treatments (with interaction term). A significant interaction would indicate that sexual and asexual species have different fundamental feeding niches. We found a significant interaction for the pair *T. californicum* - *T. shepardi*,however this was only the case for survival, and only a trend for weight gain (Table 1). We also found a significant interaction for the pair *T. poppensis* - *T. douglasi*, but only for weight gain, not survival (Table. 1). In addition, we found a marginally non-significant interaction for weight gain in the species pair *T. podura* - *T. genevievae* (Table. 1). These results suggest that in two or three species pairs, asexuals and sexuals may have diverged in their fundamental niches.

**Table 1.** Effect of experimental feeding treatments and reproductive mode on survival and weight gain of insects.

| *Timema* species pair | Factors tested in the statistical models | Survival | Weight gain |
| --- | --- | --- | --- |
| **Pair2:** | [Reproductive mode] | 1.1x10<sup>-05</sup> *** | F<sub>(1, 34)</sub> = 3.9, p = 0.054 • |
| *T. cristinae* / | [Feeding treatment] | 2.9x10<sup>-09</sup> *** | F<sub>(5, 34)</sub> = 14.8, p = 10.0x10<sup>-08</sup> *** |
| *T. monikensis* | [Reproductive mode: Feeding treatment] interaction | 0.59 | F<sub>(2, 34)</sub> = 3.9, p = 0.222 |
| **Pair3:** | [Reproductive mode] | 0.33 | F<sub>(1, 107)</sub> = 4.9, p = 0.03 * |
| *T. poppensis* / | [Feeding treatment] | 0.20 | F<sub>(6, 107)</sub> = 13.1, p = 4.6x10<sup>-11</sup> *** |
| *T. douglasi* | [Reproductive mode: Feeding treatment] interaction | 0.44 | F<sub>(6, 107)</sub> = 5.5, p = 5.4x10<sup>-05</sup> *** |
| **Pair4:** | [Reproductive mode] | 0.009 *** | F<sub>(1, 71)</sub> = 13.7, p = 0.0004 *** |
| *T. californicum* / | [Feeding treatment] | 4.8x10<sup>-05</sup> *** | F<sub>(6, 71)</sub> = 19.4, p = 2.9x10<sup>-13</sup> *** |
| *T. shepardi* | [Reproductive mode: Feeding treatment] interaction | 0.0009 *** | F<sub>(6, 71)</sub> = 1.9, p = 0.09 • |
| **Pair5:** | [Reproductive mode] | 0.0004 *** | F<sub>(1, 80)</sub> = 4.4, p = 0.04 * |
| *T. podura* / | [Feeding treatment] | 6.4x10<sup>-19</sup> *** | F<sub>(6, 80)</sub> = 22.1, p = 3.5x10<sup>-15</sup> *** |
| *T. genevieveae* | [Reproductive mode: Feeding treatment] interaction | 0.35 | F<sub>(5, 80)</sub> = 2.1, p = 0.08 • |

## DISCUSSION

We investigated if sexual and asexual stick insect species and populations differ in their realized feeding niches and how such differences come about. We find that *Timema* asexuals generally feature smaller realized feeding niches than their sexual counterparts. Specifically, in four out of five sexual-asexual *Timema* species pairs, sexuals use about twice as many plants as asexuals in nature. In the fifth species pair, *T. poppensis/T. douglasi*, sexuals and asexuals use the same number of host plants. This species pair is likely an exception to the general pattern in *Timema*because of their ability to use the hostplant redwood. We have shown in a previous study that sexual *Timema* species adapted to this specific host plant are ecologically highly specialized, perhaps because of reduced biotic pressures on redwood (Larose et al. 2018). This high level of ecological specialization in the sexual makes further specialization in the related asexual relatively unlikely.

To develop insights into how the narrower realized niches of asexual versus sexual *Timema* species come about, we quantified the size of their fundamental feeding niches. This allowed us to test if the size of the fundamental niche constrains the size of the realized niche, i.e., whether the reduced realized niche size in asexuals results from a reduced intrinsic ability to use different host plants. Fundamental feeding niche size varied significantly among all *Timema* species, however there was no overall difference between reproductive modes. Fundamental niche size therefore does not explain why sexuals have broader realized niches than asexuals in *Timema*. Specifically, in two species pairs the estimated fundamental niche size was very similar for sexuals and asexuals. In the other two pairs, the fundamental niche differed between sexuals and asexuals, however in opposite directions; In one species pair (*T. cristinae/T. monikensis*) the asexual species had a narrower fundamental niche than the sexual one, while in the other (*T. podura/T. genevievae*) the asexual species had a broader fundamental niche than the sexual one. The latter case is particularly interesting because *T. genevievae* is a very old asexual lineage (~1.5-2 myr) and the oldest asexual *Timema* known (Schwander *et al*. 2011). The broad fundamental feeding niche in *T. genevievae* is consistent with predictions from the *General Purpose Genotype* (GPG) theory, which posits that clones with broad environmental tolerances (i.e., broad fundamental niches) should be selectively favored as such clones would be characterized by low variance in fitness across environments (Lynch 1984; Fig. 1B). General purpose genotypes are also believed to contribute to the persistence of one of the oldest known asexual species, the darwinulid ostracod *Darwinula stevensoni*, which has probably existed as an obligate asexual for 25 million years (Straub 1952). It shows almost no morphological (Rossetti and Martens 1998) or genetic (Schön et al. 1998) variability, yet it is a very common and cosmopolitan species (Griffiths and Butlin 1994) with broad tolerances for salinity and temperature (Van Doninck et al. 2002).

In contrast to the old asexual *T. genevievae*, our findings in the youngest studied *Timema* asexual, *T. monikensis*, are consistent with the *Frozen niche variation* model (FNV). This model suggests that the phenotypic distribution (i.e., fundamental niche) of a young, recently derived asexual lineage will be narrower than that of its genetically variable sexual ancestor (Vrijenhoek 1984; Fig. 1A). Indeed, *T. monikensis* is the only studied asexual that features a narrower fundamental niche than its sexual relative *T. cristinae* (Figs. 3B; Fig. S1A).

Given that asexual *Timema* do not generally have narrower fundamental niches than sexual *Timema*, the narrow realized niches in asexuals are likely a consequence of biotic interactions that affect niche size in asexuals more strongly than in sexuals. A likely biotic factor affecting realized niches in *Timema* is selection imposed by predators (e.g., Sandoval 1994a, 1994b, Nosil et al. 2003, Nosil 2004). Several *Timema* species feature a natural color polymorphism conferring crypsis on different host plants (Sandoval 1994a; Sandoval 1994b) and we therefore tested for links between color polymorphism, realized niche size and reproductive mode in *Timema*. The sister species *T. douglasi* and *T. poppensis* do not feature any color polymorphism, but in the four remaining species pairs, intra-population color polymorphism is always higher in the sexual than asexual species. However, the level of polymorphism was only correlated to the size of the realized niche at the species level, not at the population level. Nevertheless, this higher degree of color polymorphism in sexuals may allow for reduced predation rates on a larger number of plants relative to asexuals, potentially explaining the narrower realized niche size in asexual species.

In conclusion, we provide the first comparative study of realized and fundamental niches in replicated asexual-sexual species pairs. We found that sexual *Timema* species have a larger realized niche than asexual ones, but this difference is not explained by a similar difference in fundamental niche size. Thus, the smaller realized niches in asexuals are likely a consequence of biotic interactions that constrain asexuals more strongly than sexuals. Verifying potential links between population-level color polymorphism, realized feeding niche size and biotic interactions (especially predation and competition) will be a challenge for future studies. Finally, our finding that the oldest asexual *Timema* lineage is more generalist than it sexual relative could help explain its unusually long maintenance in the absence of sex.

## Acknowledgments

We thank Armand Yazdani and Ian S. Ford for their help in the field, and Giacomo Bernardi at UC Santa-Cruz for labspace. This study was supported by grants PP00P3_139013 and PP00P3_170627of the Swiss FNS to TS.

## SUPPORTING INFORMATION

**Table S1.** Color morphs of ten *Timema* species

| *Timema* species | Color morphs<sup>1</sup> |  |  |  |  |  |  |  |  |  |  |  |  |  |
| --- | --- | --- | --- | --- | --- | --- | --- | --- | --- | --- | --- | --- | --- | --- |
|  | b | bl | br | db | dg | g | gr | oc | ol | sg | sgp | r | y | w |
| *T. bartmani* |  |  | x |  |  | x | x |  | x | x | x |  |  |  |
| *T. tahoe* |  |  | x |  |  | x |  |  |  | x | x |  |  |  |
| *T. cristinae* | x |  |  | x |  | x | x | x |  | x |  | x | x |  |
| *T. monikensis* | x |  |  | x |  | x |  |  |  |  |  |  | x |  |
| *T. poppensis* |  |  |  |  |  | x |  |  |  |  |  |  | (x) |  |
| *T. douglasi* |  |  |  |  |  | x |  |  |  |  |  |  | (x) |  |
| *T. californicum* | x | (x) |  |  |  | x | x |  |  |  |  | x | x | (x) |
| *T. shepardii* |  |  |  |  |  | x |  |  |  |  |  |  | (x) |  |
| *T. podura* | x |  | x | x | x | x |  |  | x |  |  |  |  |  |
| *T. genevieveae* |  |  |  |  | x |  |  |  |  |  |  |  |  |  |

<sup>1</sup>b, beige; bl, blue ; br, brown, db, dark brown, dg, dark grey; g, green; gr, grey; oc, ochre; ol, olive; sg, striped green; sgp, striped green with pink head; r, red; y, yellow; w, white
A cross indicates that a specific morph has been observed in a given species. A cross in brackets indicates that the morph was observed at a frequency of less than 0.1% in a given species, all populations combined.

**Table S2.**
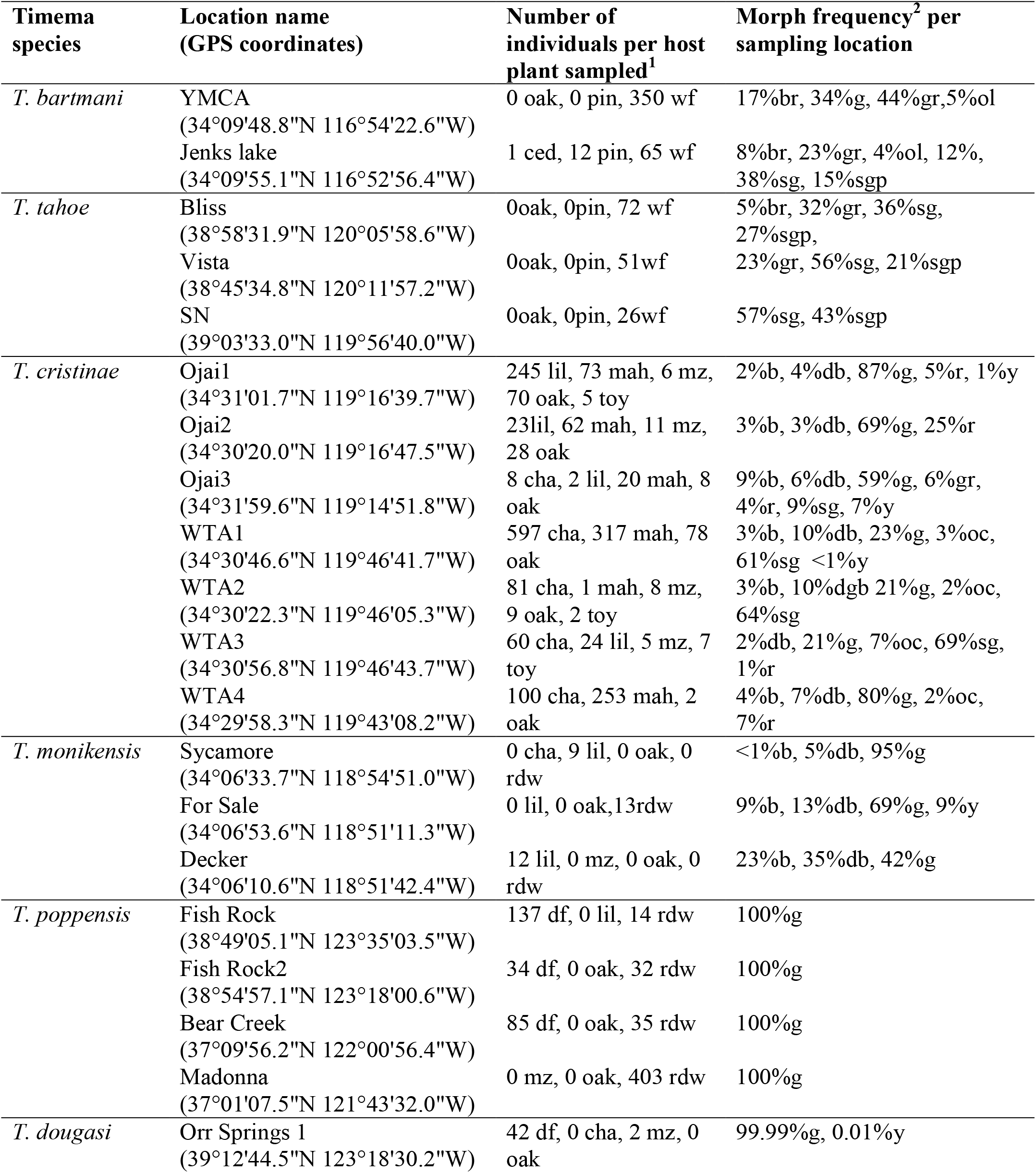
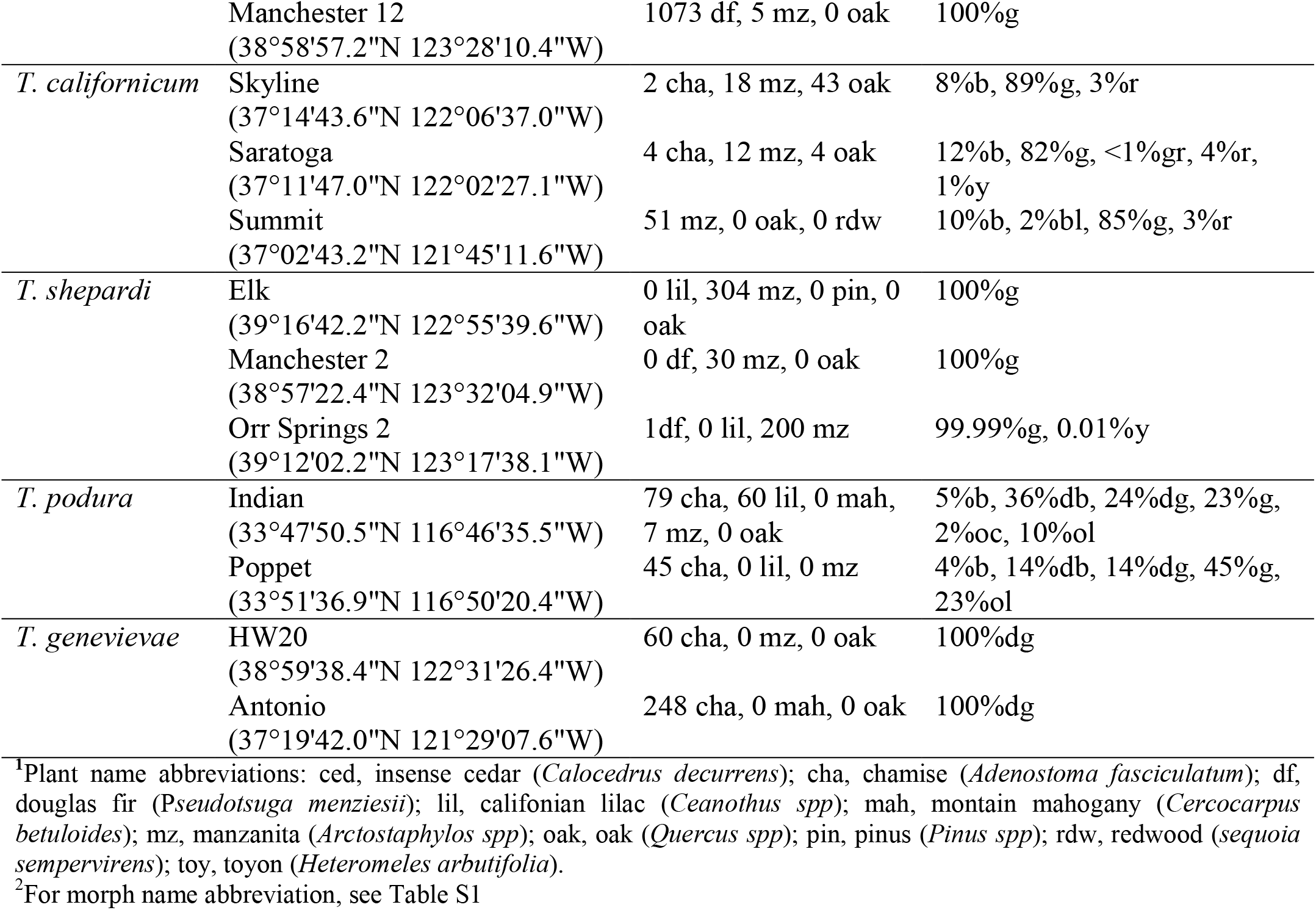
*Timema* populations sampled. Number of individuals refers to the total number of individuals sampled in these locations on different host plants. We only selected locations in which at least three plants from the known host plant set were present.

**Table S3.**
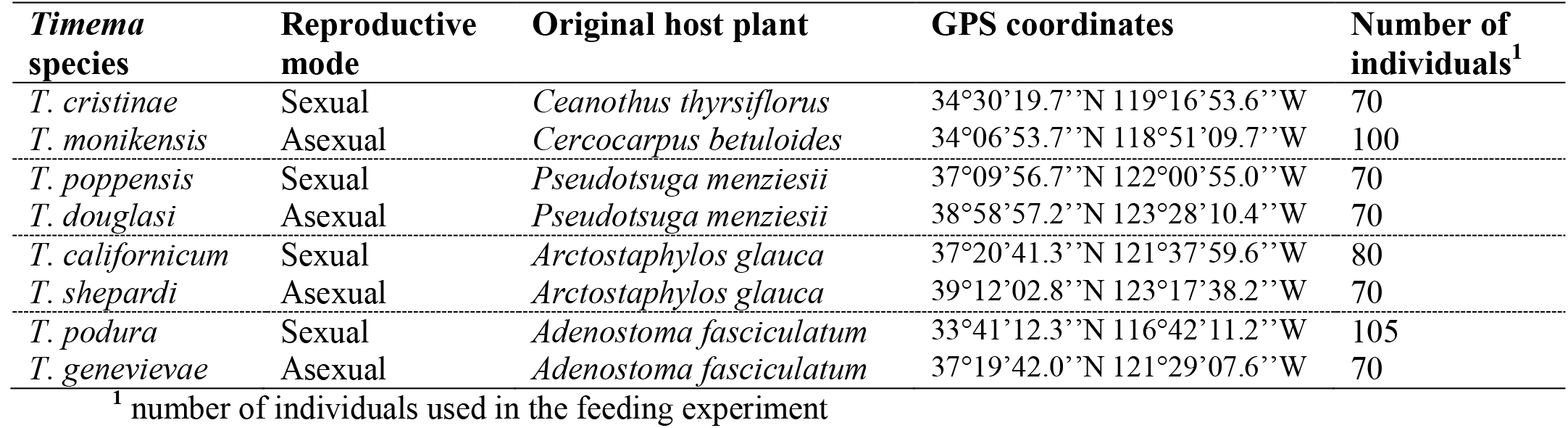
Overview of the *Timema* ssp used for the study of the fundamental feeding niche

| ***Timema***<br>**species** | **Reproductive**<br>**mode** | **Original host plant** | **GPS coordinates** | **Number of**<br>**individuals<sup>1</sup>** |
| --- | --- | --- | --- | --- |
| *T. cristinae* | Sexual | *Ceanothus thyrsiflorus* | 34°30'19.7''N 119°16'53.6''W | 70 |
| *T. monikensis* | Asexual | *Cercocarpus betuloides* | 34°06'53.7''N 118°51'09.7''W | 100 |
| *T. poppensis* | Sexual | *Pseudotsuga menziesii* | 37°09'56.7''N 122°00'55.0''W | 70 |
| *T. douglasi* | Asexual | *Pseudotsuga menziesii* | 38°58'57.2''N 123°28'10.4''W | 70 |
| *T. californicum* | Sexual | *Arctostaphylos glauca* | 37°20'41.3''N 121°37'59.6''W | 80 |
| *T. shepardii* | Asexual | *Arctostaphylos glauca* | 39°12'02.8''N 123°17'38.2''W | 70 |
| *T. podura* | Sexual | *Adenostoma fasciculatum* | 33°41'12.3''N 116°42'11.2''W | 105 |
| *T. genevieveae* | Asexual | *Adenostoma fasciculatum* | 37°19'42.0''N 121°29'07.6''W | 70 |

<sup>1</sup> number of individuals used in the feeding experiment

**Figure S1.**
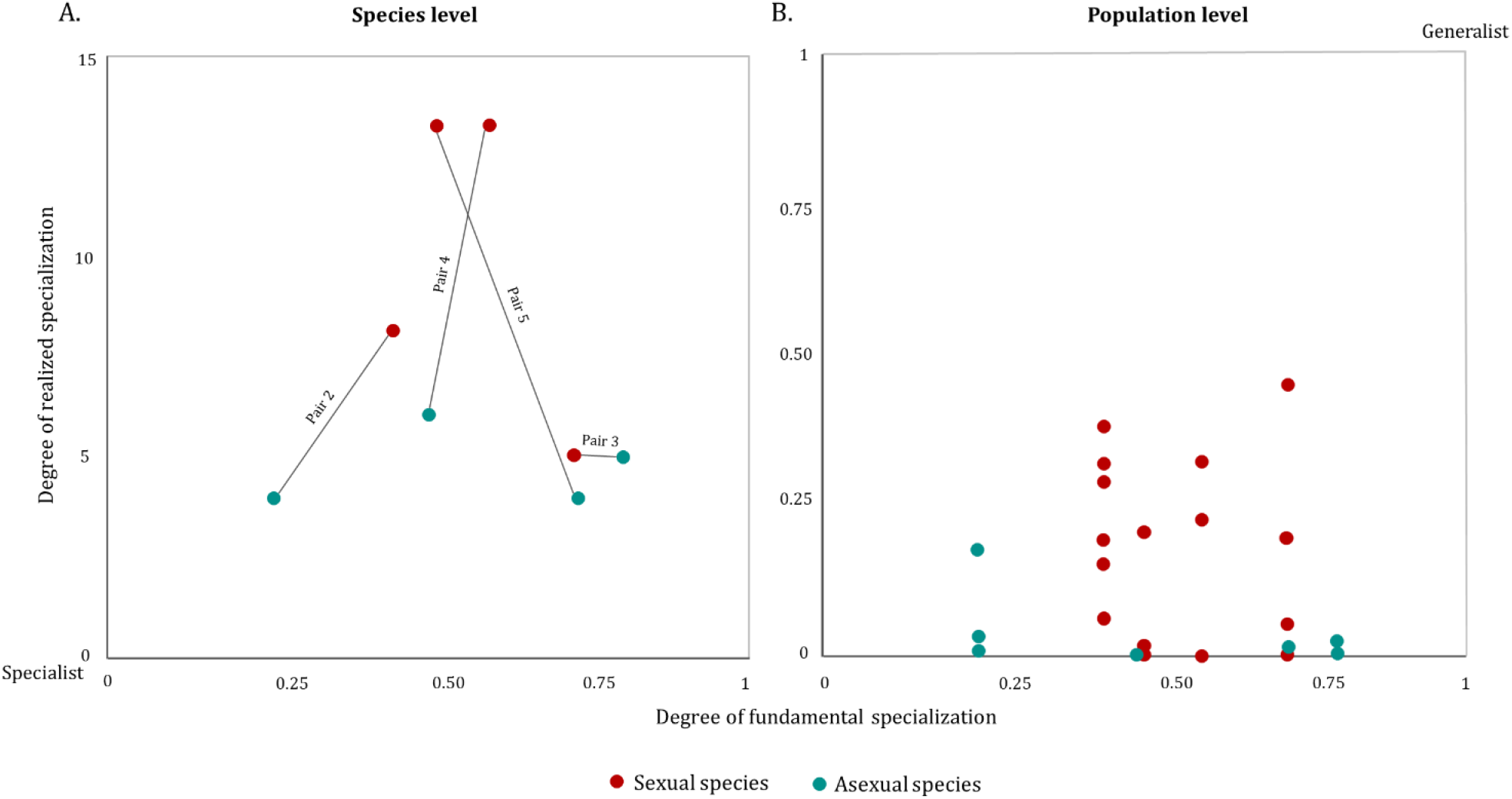
Realized and fundamental feeding niche breadths of sexual and asexual stick insects are not correlated. Shown is the specificity index Tau (calculated from the weight gain of insects) as a function of the realized feeding niche at the species level **(A)** or at the population level **(B)**. For species pair numbers, see Fig. 2 in the main text.

**Figure S2.**
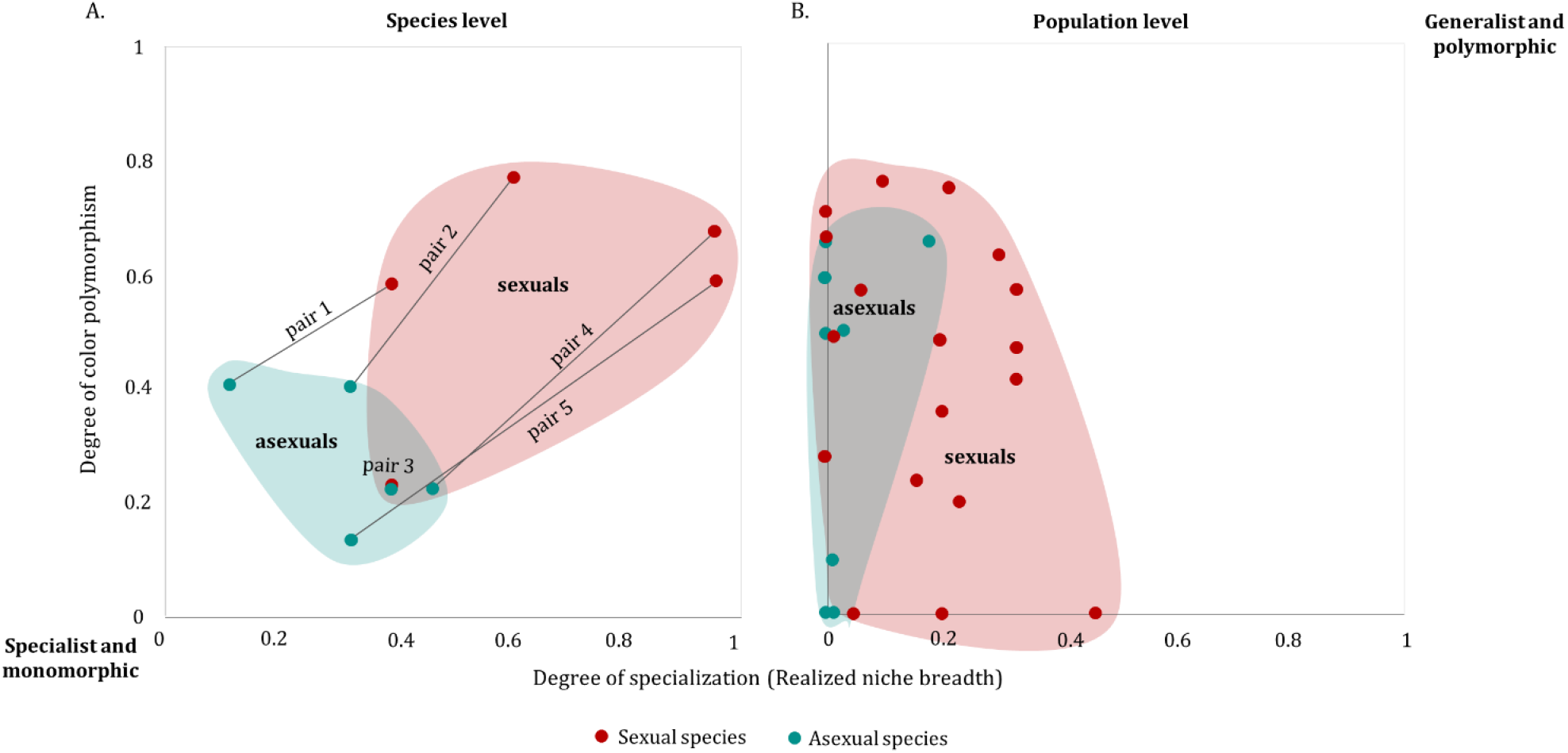
Correlation between color polymorphism and realized feeding niche breadth of Timema at the species (A) and at the population **(B)** levels. At the species level (A), the polymorphism levels and realized feeding niche sizes are estimated from a count of the different color morphs and of the known host plants in each species respectively. At the population level (B), the polymorphism level is estimated using the inverse Simpson diversity index, and the realized feeding niche size is estimated using the Tau index. In this case, 0 corresponds to specialism and monomorphism, and 1 corresponds to generalism and extreme polymorphism.

